## Supplementary material for "Multi-omics evaluation of cell lines as models for metastatic prostate cancer": Supp Figures

Liu et al.

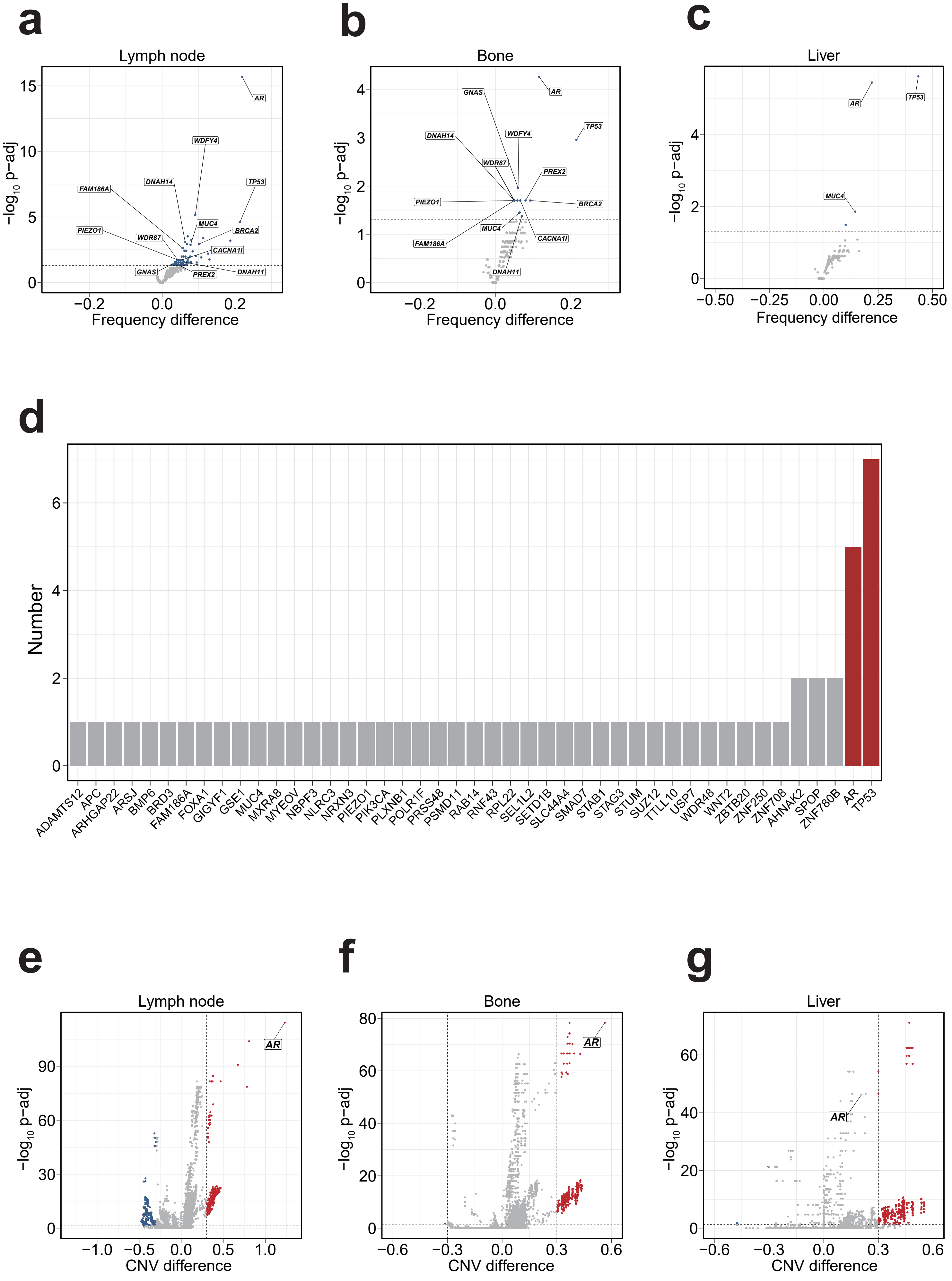


Supplementary Figure 1

**Supplementary Figure 1:**

(a-c) Metastasis-site-specific comparisons of gene mutation frequencies between SU2C and TCGA samples. Each point represents a gene; the x-axis denotes the difference in mutation frequency (SU2C-TCGA), and the y-axis shows statistical significance. The horizontal dashed line indicates an adjusted *P*-value cutoff of 0.05 and the differentially mutated genes are labeled.

(d) Ranking 45 genes based on the number of carried mutation hotspots.

(e-g) Metastasis-site-specific comparisons of gene copy number variation (CNV) profiles between SU2C and TCGA samples. Each point represents a gene; the x-axis denotes the difference in median CNV values (SU2C – TCGA), and the y-axis shows statistical significance. Dashed vertical lines represent CNV difference thresholds (±0.3), and the horizontal dashed line indicates an adjusted *P*-value cutoff of 0.05.

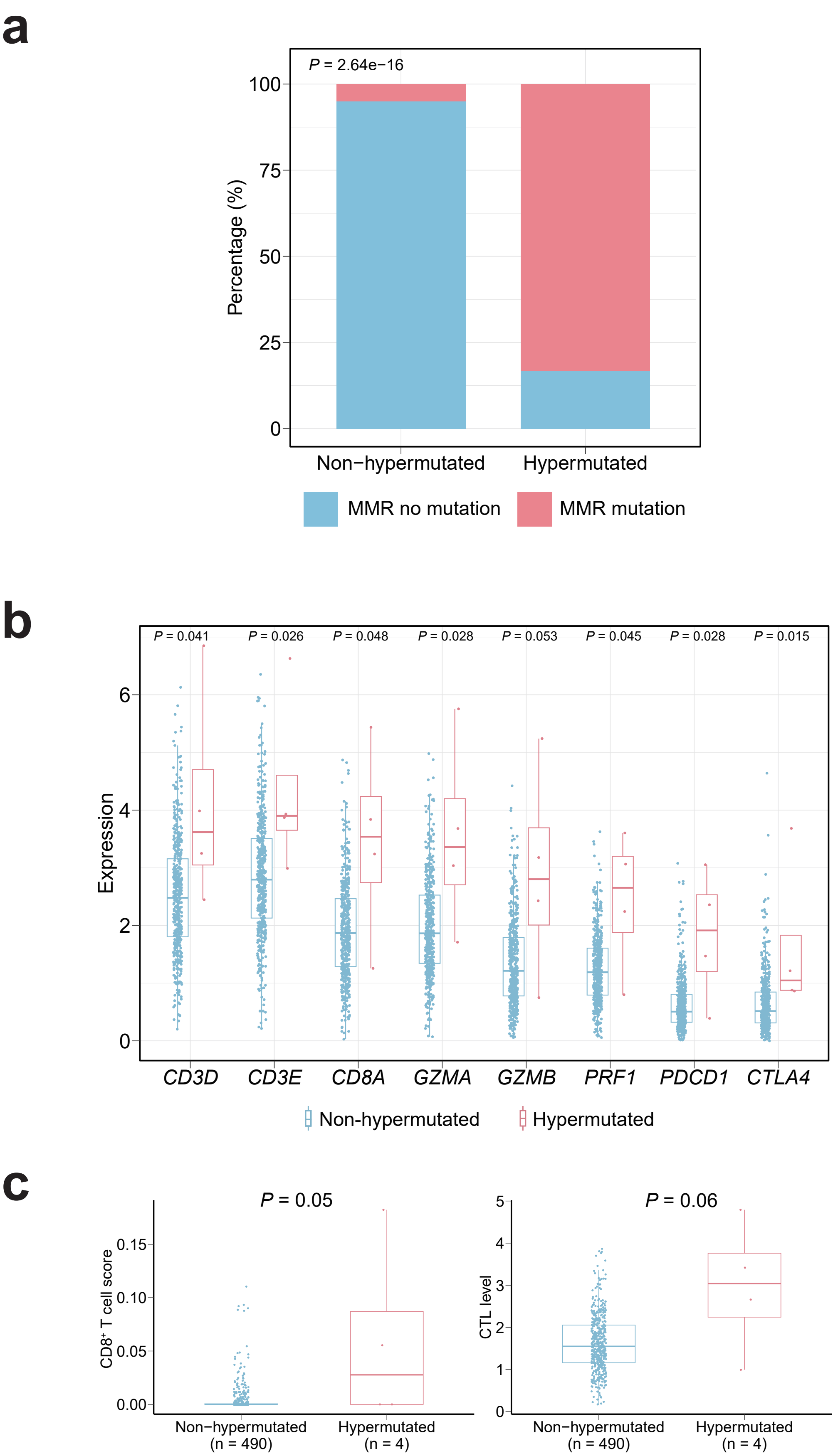


**Supplementary Figure 2**

**Supplementary Figure 2:**

(a) Comparison of mutation frequencies of MMR genes between hypermutated and non-hypermutated samples. *P*-value was calculated using Fisher's exact test.

(b) Expression of immune-related genes (*CD3D*, *CD3E*, *CD8A*, *GZMA*, *GZMB*, *PRF1*, *PDCD1*, and *CTLA4*) in hypermutated (n=4) and non-hypermutated (n = 490) TCGA samples. In each box, the central line represents the median value and the bounds represent the 25th and 75th percentiles (interquartile range). The whiskers encompass 1.5 times the interquartile range.

(c) Comparison of $CD8^+$ T cell scores estimated by the xCell algorithm (left) and cytotoxic T lymphocyte (CTL) levels estimated by the TIDE algorithm (right). In each box, the central line represents the median value and the bounds represent the 25th and 75th percentiles (interquartile range). The whiskers encompass 1.5 times the interquartile range.

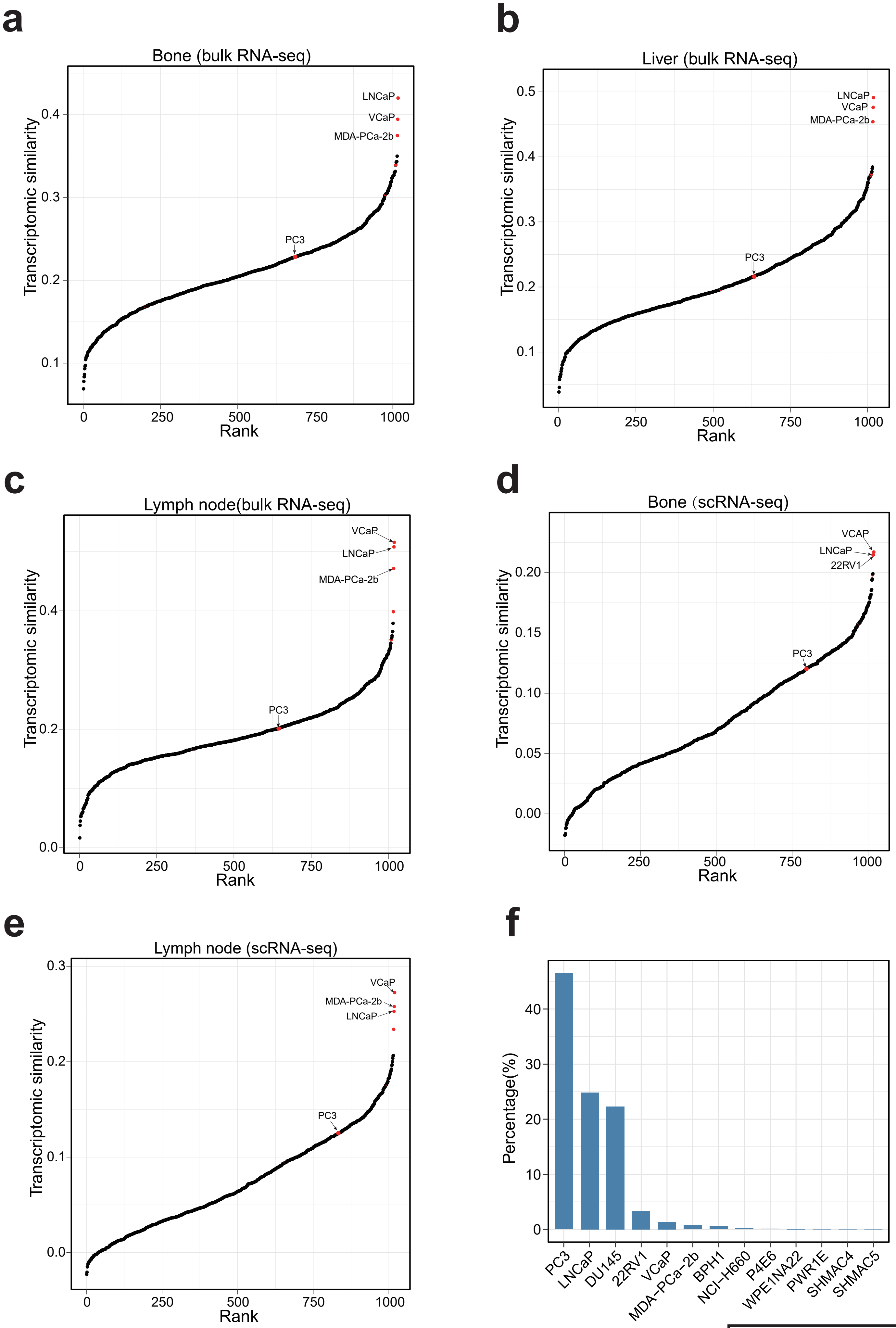


**Supplementary Figure 3**

**Supplementary Figure 3:**

(a-c) Ranking 1,019 CCLE cell lines based on their transcriptomic similarity to MET500 prostate cancer samples from bone (a), liver (b), and lymph-node (c) metastases. Each dot represents a CCLE cell line, and the prostate cancer cell lines are highlighted in red.

(d-e) Ranking 1,019 CCLE cell lines based on their transcriptomic similarity to the malignant cells from (d) bone and (e) lymph-node metastases in a single-cell RNA-seq dataset. Each dot represents a CCLE cell line, and the prostate cancer cell lines are highlighted in red.

(f) Ranking CCLE prostate cancer cell lines based on PubMed citation frequency.

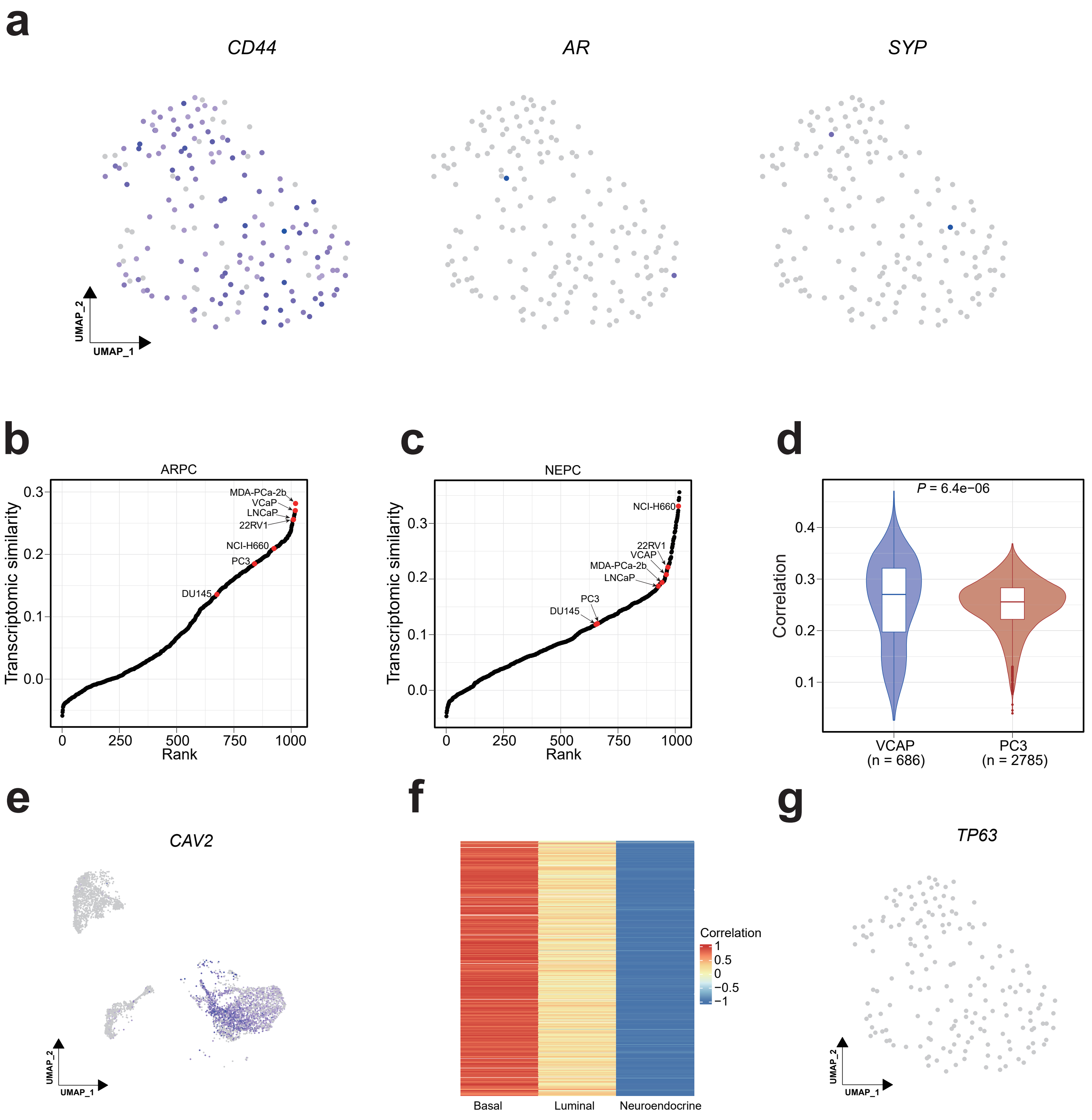


**Supplementary Figure 4**

**Supplementary Figure 4:**

(a) Expression of *CD44*, *AR*, *SYP* in PC3 single cells. Color represents expression level, from gray (low) to dark blue (high).

(b-c) Ranking 1,019 CCLE cell lines based on their transcriptomic similarity to the malignant cells of (b) ARPC and (c) NEPC subtypes. Each dot represents a CCLE cell line, and the prostate cancer cell lines are marked highlighted in red.

(d) Transcriptomic correlation between VCaP and ARPC malignant cells is significantly higher than that between PC3 and MSPC malignant cells. In each box, the central line represents the median value and the bounds represent the 25th and 75th percentiles (interquartile range). The whiskers encompass 1.5 times the interquartile range. Outliers are shown as individual points.

(e) Expression of *CAV2* in malignant cells from the CRPC scRNA-seq dataset. Color represents expression level, from gray (low) to dark blue (high).

(f) Heatmap showing the transcriptomic correlation between MSPC malignant cells and three normal prostate epithelial cell types.

(g) Expression of *TP63* in PC3 single cells. All cells show no detectable *TP63* expression (gray).

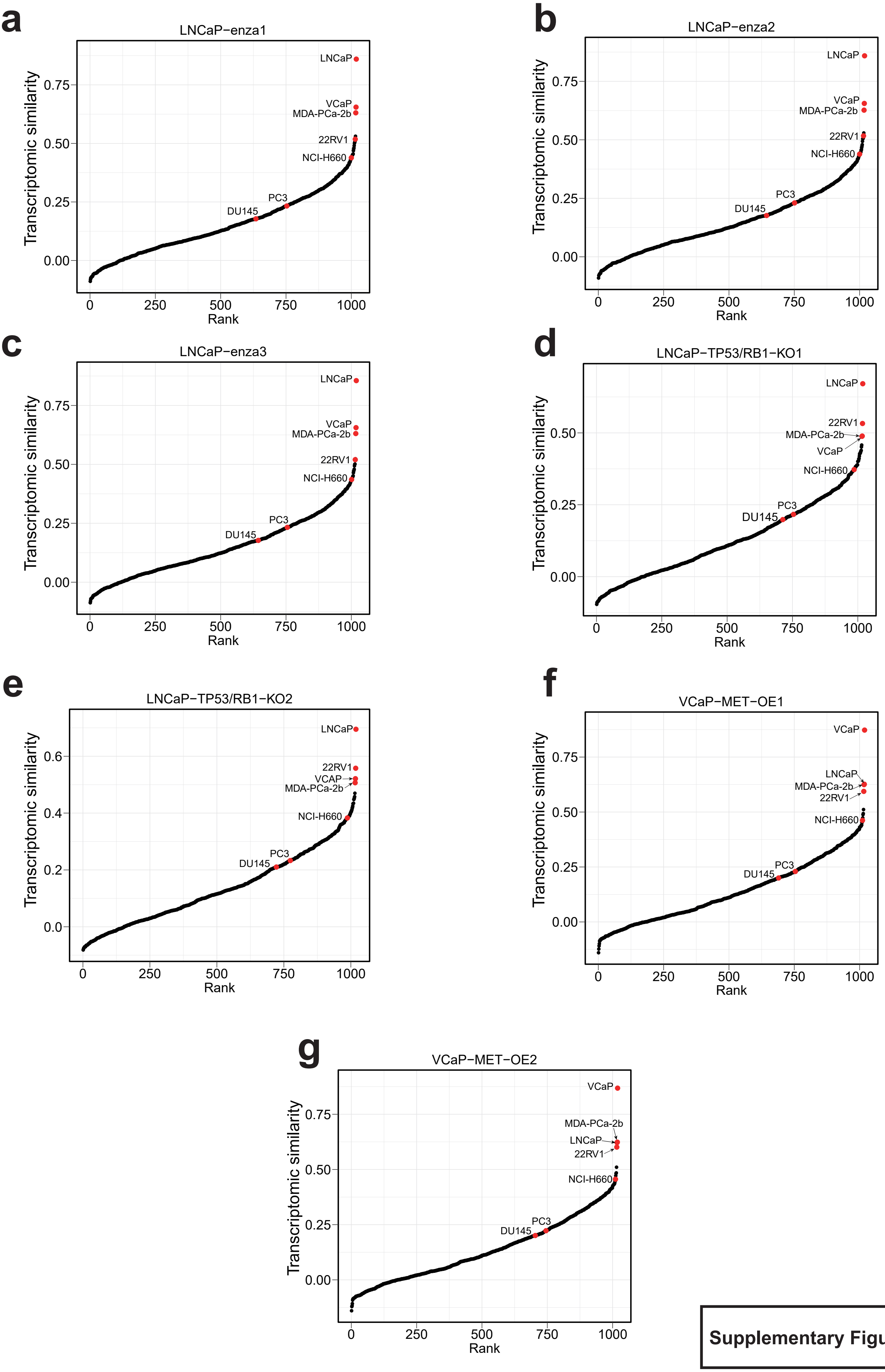


**Supplementary Figure 5**

**Supplementary Figure 5:**

(a-c) Ranking 1,019 CCLE cell lines based on their transcriptomic similarity to engineered LNCaP cell lines (treated with enzalutamide). Each dot represents a CCLE cell line, and the prostate cancer cell lines are highlighted in red.

(d-e) Ranking 1,019 CCLE cell lines based on their transcriptomic similarity to engineered LNCaP cell lines (*TP53*/*RB1* double knockout). Each dot represents a CCLE cell line, and the prostate cancer cell lines are highlighted in red.

(f-g) Ranking 1,019 CCLE cell lines based on their transcriptomic similarity to engineered VCaP cell line (*MET* overexpression). Each dot represents a CCLE cell line, and the prostate cancer cell lines are highlighted in red.
